# Computer Vision Methods for Spatial Transcriptomics: A Survey

**DOI:** 10.1101/2025.10.13.682148

**Authors:** Junchao Zhu, Ruining Deng, Junlin Guo, Tianyuan Yao, Siqi Lu, Chongyu Qu, Juming Xiong, Yanfan Zhu, Zhengyi Lu, Yuechen Yang, Marilyn Lionts, Yucheng Tang, Daguang Xu, Yu Wang, Shilin Zhao, Haichun Yang, Yuankai Huo

## Abstract

Spatial transcriptomics (ST) enables the simultaneous measurement of gene expression and spatial localization within tissue sections, providing unprecedented opportunities to dissect tissue architecture and functional organization. As a relatively new omics technology, bioinformatics has driven much of the innovation in ST. However, within these frameworks, “spatial” information is often reduced to locations and relationships between molecular profiles, without fully leveraging the wealth of submicron morphological detail and histological knowledge available. Advances in computer vision–based artificial intelligence (AI) are opening exciting new avenues beyond conventional bioinformatics approaches by modeling complex histological patterns and linking morphology to molecular states. More excitingly, they bring fresh perspectives to potentially address key limitations of ST, including its high cost, limited clinical applicability, and reliance on twodimensional (2D) analysis of inherently three-dimensional (3D) tissues. For instance, models that predict ST directly from histology images enable “virtual sequencing,” drastically reducing costs while integrating morphological insights from pathology with molecular biomarkers, thus accelerating clinical translation. Moreover, computer vision techniques can reconstruct pixel-aligned 3D tissue models, overcoming the technical barriers of 2D acquisition and advancing 3D spatial omics analytics. In this paper, we present the first systematic survey of computer vision AI models for ST analytics, categorizing approaches across architectures, learning paradigms, tasks, and datasets, and tracing their technological evolution. We highlight key challenges and future directions, offering a panoramic perspective on vision-driven ST and its potential to transform both basic research and clinical practice. The curated collection of vision-driven spatial transcriptomics papers is available at https://github.com/hrlblab/computer_vision_spatial_omics.

## I. Introduction

As a rapidly advancing multi-omics approach, spatial transcriptomics (ST) enables the simultaneous capture of gene expression and spatial localization within tissue sections, offering unprecedented insights into cellular states, tissue architecture, and their functional organization [4], [7]. It has shown great potential in diverse applications such as tumor microenvironment analysis, neuroscience, developmental biology, and clinical pathology [5], [59].

However, as a relatively new omics technology, ST still relies heavily on bioinformatics methods for data analysis and model construction. Existing studies mainly focus on processing molecular expression matrices, modeling spatial adjacency relationships, and constructing gene co-expression networks [8], [76], [82]. Although these methods have achieved remarkable progress at the expression level, they often reduce spatial information to mere coordinates and relational graphs, overlooking the submicron-scale morphological details and rich histological knowledge embedded in tissue sections [26]. In fact, histological images contain highly intricate structural signals, from nuclear morphology to tissue-level organization, that are tightly correlated with gene expression [3], yet have long been underestimated in traditional bioinformatics analyses.

In recent years, rapid advances in computer vision and artificial intelligence (AI) [20], [27] have injected new momentum into ST analytics [11], [74], [119]. Deep learning models can uncover visual patterns and capture nonlinear mappings between tissue structure and molecular states [26], [100], [106], realizing virtual sequencing. This paradigm shift signifies the transition for clinical pathology [40], [108].

Furthermore, computer vision techniques are also driving the development of three-dimensional spatial omics [104], [119]. By registering, reconstructing, and pixel-aligning consecutive sections, researchers can obtain comprehensive 3D molecular maps of tissues, thereby overcoming the limitations of 2D data acquisition [46]. Such cross-dimensional integration enables more holistic and systematic investigations of tissue development, disease progression, and tumor microenvironments, while laying the foundation for future multimodal omics integration.

Despite the remarkable potential of vision-driven ST models, the field remains in a rapidly evolving stage, lacking systematic summarization and directional guidance. This paper aims to provide a comprehensive and up-to-date overview of vision-driven approaches for spatial transcriptomics, with a focus on representative models and techniques proposed between 2020 and 2025. This period has witnessed a rapid evolution of AI-based methods in the ST domain. While early studies established the feasibility of predicting gene expression from WSIs, most existing reviews have centered on the evolution of ST platforms, with limited attention to AI-driven image-based modeling. Therefore, this survey specifically highlights the methodological innovations and unique contributions of vision-driven ST in the past five years, aiming to synthesize key developments, identify remaining challenges, and offer perspectives for future research.

Figure 1 presents a chronological overview of 46 representative vision-driven ST models published between 2020 and 2025, illustrating the field’s evolution from early-stage explorations to its recent rapid expansion. To enhance clarity and practical utility, we further categorize these models based on the spatial resolution level targeted by each method and the type of downstream tasks they address. This structure helps clarify which models have been applied to general ST challenges versus more specific biological or clinical applications. It is important to note that the inclusion of a model in this timeline reflects existing validation, but does not limit the potential of other architectures. In fact, advanced paradigms from computer vision and natural language processing are increasingly being adapted to ST research [17], [34], [57].

**Fig. 1:**
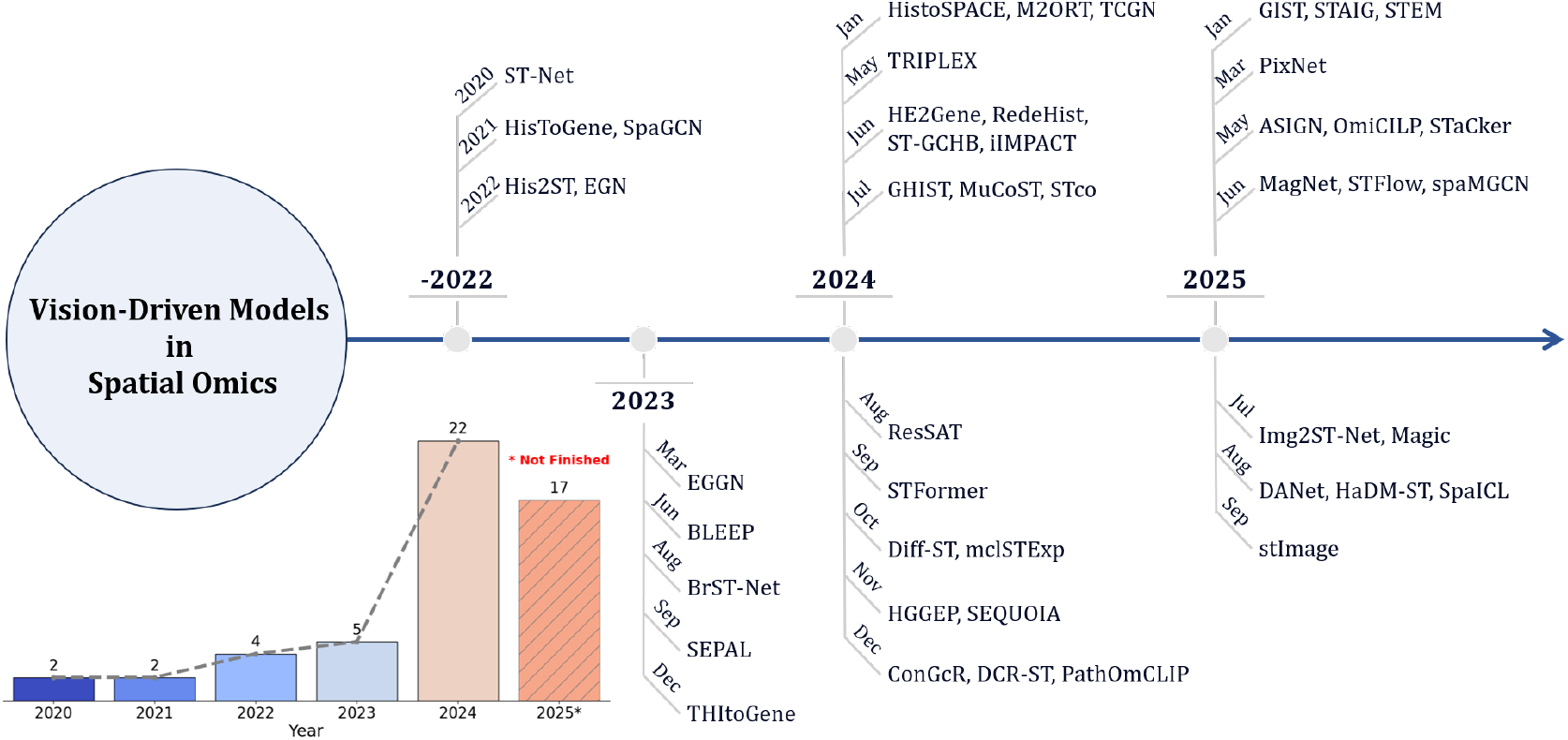
Overview of some well-known vision-driven models for spatial transcriptomics from 2020 to 2025 September. of ST research and offers new molecular-level insights

Our contributions include:

1. The first comprehensive review of existing vision-driven ST models, covering methodological background, core architectures, and downstream applications. We summarize recent developments in a hierarchical and structured manner, highlighting the evolution of the field and the connections across different research directions.
2. Categorization and comparative analysis of models based on their training paradigms (Table III) and real-world application scenarios (Table I). We focus on architectural designs, training strategies, data requirements, and task suitability, aiming to provide a clear understanding of each model’s strengths, limitations, and practical scope.
3. Discussion of open challenges and future directions in vision-driven ST. We identify key bottlenecks, emerging trends, and unresolved questions, and offer insights into potential avenues for future research.

**TABLE I:**
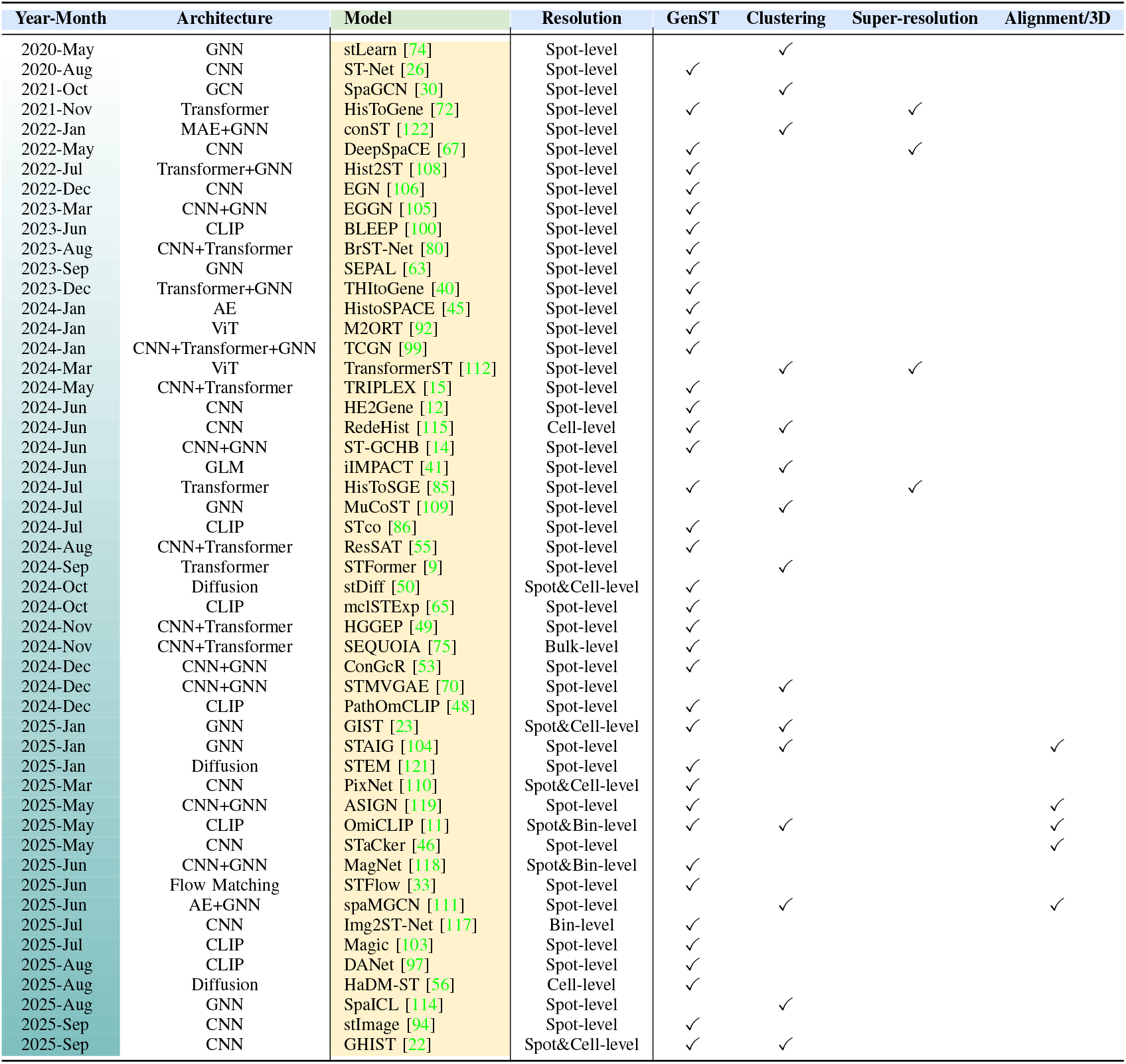
Landscape of vision-driven models and downstream tasks from May 2020 to Sep 2025. For each model, we summarize the backbone architecture, output resolution, and reported tasks. Checkmarks denote tasks that were quantitatively evaluated.

## II. Data Landscape

### A. Spatial Transcriptomics Data Acquisition

ST maps gene expression to precise locations within intact tissue architecture [61]. By retaining the spatial coordinates of each gene’s expression within tissue sections, ST enables in situ analyses and allows researchers to investigate the interactions between cellular functional states and their microenvironments in situ [96], [116]. It captures gene expression patterns directly within the spatial context of tissue slices, thereby aligning morphological and molecular evidence on the same slide [51], [81].

As illustrated in Figure 2, ST technologies span a wide range of spatial resolutions, from lower-resolution platforms like traditional Visium ( 55 µm spot size), to high-resolution and subcellular-scale imaging platforms such as Xenium or CosMx. Broadly, ST technologies can be categorized into two main types [68], including in Situ Sequencing (ISS)-based approaches [88], [89], and in situ hybridization-based (ISH) methods [21], [98]. The ISS uses spatially barcoded arrays to capture mRNA for high-throughput sequencing, enabling broad tissue coverage, but is limited by spot size, which may mix signals from multiple cells. The ISH employs fluorescent probes for in situ labeling and iterative imaging, offering single-cell or subcellular resolution, ideal for studying cellular heterogeneity, though it is more costly, limited in gene coverage, and experimentally complex. Each platform presents distinct strengths and limitations in terms of spatial resolution, gene detection throughput, quantification accuracy, and compatibility with different tissue types [13], [52], [93].

**Fig. 2:**
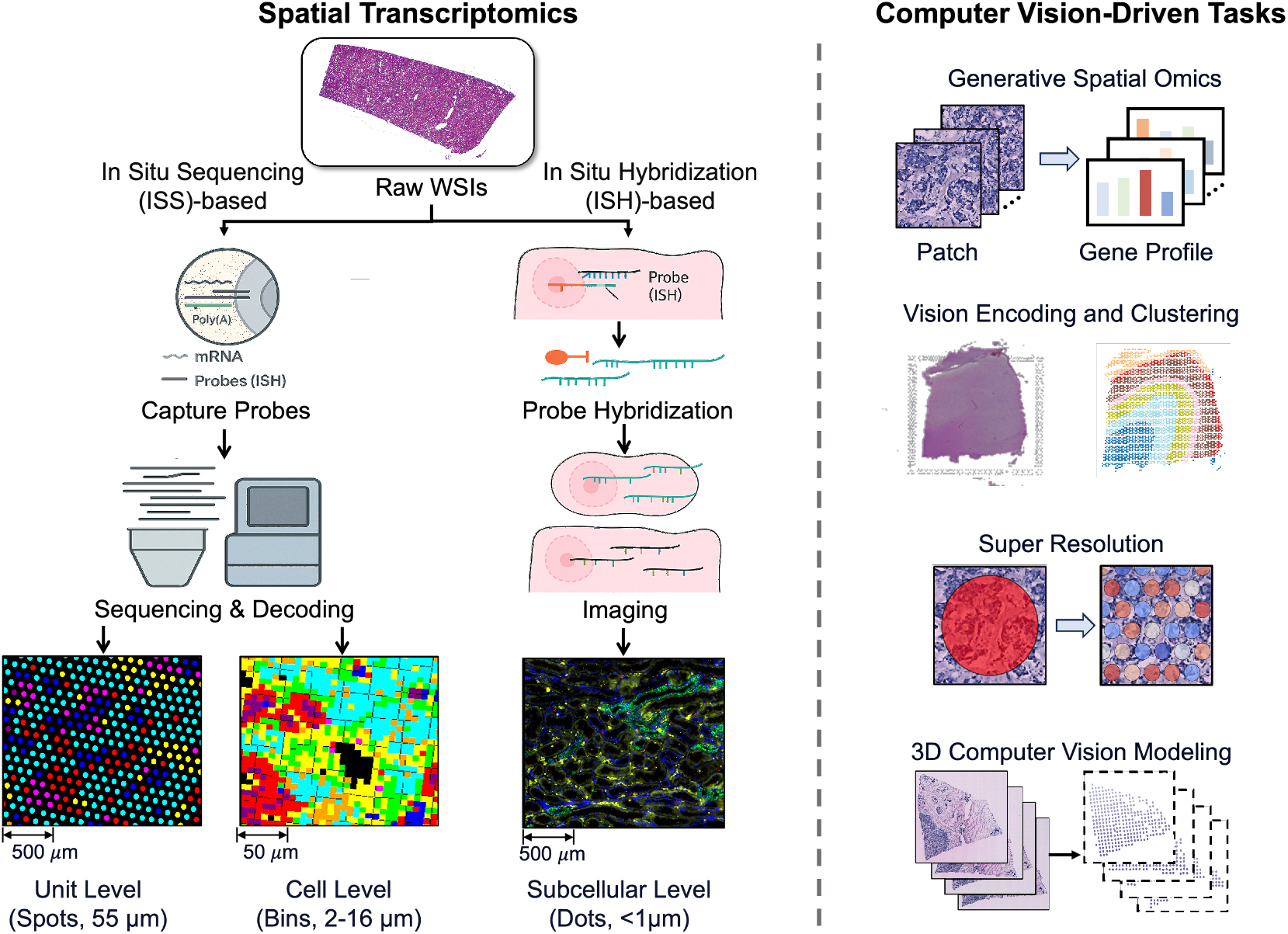
Spatial Transcriptomics Data Acquisition and Computer Vision Methods. Left: Two acquisition families. (a) In Situ Sequencing (ISS)-based capture producing 55 µm spot-level maps and high-resolution binned maps (2/8/16 µm). (b) In-situ hybridization yielding subcellular readouts (<1 µm). Right: Representative downstream tasks enabled by histology and ST, including (1) Generative Spatial Omics, (2) Vision Feature Encoding and Clustering, (3) Super Resolution, (4) 3D Computer Vision Modeling.

With the development of high-resolution platforms and multi-omics integration, ST has been broadly applied in tumor microenvironment studies, neural development, immune profiling, pathology, and integrative biomedical research [16], [42], [77], facilitating the analysis of tissue functional zonation and cell–cell spatial interactions [107]. Moreover, several studies have combined ST with complementary modalities, including singlecell RNA sequencing, spatial proteomics, and digital pathology imaging, to enhance biological interpretability and predictive power [26], [66].

Despite the rapid progress of ST, data analysis still faces major challenges, including high-dimensional and sparse expression matrices, cross-platform batch effects, spatial misalignment, and the lack of standardized annotations [4], [13], [61]. A variety of strategies have been proposed [19], [83], [113] to facilitate ST analysis.

### B. Evolution of Public ST Datasets

Over the past decade, public ST datasets have undergone rapid evolution, with remarkable improvements in resolution, scale, and annotation. Table II summarizes these developments, outlining a clear trajectory: from spot-level profiling to multi-resolution analyses, from small-scale exploratory datasets to large-scale population cohorts, from single-organ studies to multi-organ atlases, and from supporting single-task analyses to enabling multi-task annotation resources.

**TABLE II:** Summary of representative spatial transcriptomics datasets used in image-to-ST studies. For each dataset, we report the release date, dataset name, profiling resolution, primary organs, number of samples and slides, total spots, and typical applications.

| Year-Mon | Dataset | Title | Resolution | Organ | Sample | Slides | Spot | Application |
| --- | --- | --- | --- | --- | --- | --- | --- | --- |
| 2016-Jul | MOB [88] | Visualization and analysis of gene expression in tissue sections by spatial transcriptomics | Spot-level | Brain | 1 | 12 | 4,147 | ST Prediction |
| 2020-Jul | cSCC [38] | Multimodal analysis of composition and spatial architecture in human squamous cell carcinoma | Spot-level | Skin | 4 | 12 | 8,671 | Alignment/3D, ST Prediction |
| 2020-Aug | Breast-ST [26] | Integrating spatial gene expression and breast tumour morphology via deep learning | Spot-level | Breast | 23 | 68 | 30,655 | Alignment/3D, ST Prediction |
| 2021-Feb | DLPFC [62] | Transcriptome-scale spatial gene expression in the human dorsolateral prefrontal cortex | Spot-level | Brain | 3 | 12 | 47,681 | Clustering, Alignment/3D, ST Prediction |
| 2021-Oct | HER2 [1] | Spatial deconvolution of HER2-positive breast cancer delineates tumor-associated cell type interactions | Spot-level | Breast | 8 | 36 | 13,620 | Alignment/3D, ST Prediction |
| 2023-Jul | Human-Kidney [47] | An atlas of healthy and injured cell states and niches in the human kidney | Spot-level | Kidney | 24 | 24 | 74,220 | ST Prediction |
| 2023-Jul | Human-CCS [43] | Spatially resolved multiomics of human cardiac niches | Spot-level | Heart | 25 | 41 | 100,539 | ST Prediction, Clustering |
| 2023-Nov | Breast-HD [69] | Visium HD enables spatial discovery in FFPE human breast cancer at single-cell scale | Bin-level | Breast | 4 | 4 | - | ST Prediction |
| 2023-Dec | BC1 [36], BC2 [36] | High resolution mapping of the tumor microenvironment using integrated single-cell, spatial and in situ analysis | Spot-level | Breast | 2 | 2 | 6,331 | ST Prediction |
| 2023-Dec | Breast-Xenium [36] | High resolution mapping of the tumor microenvironment using integrated single-cell, spatial and in situ analysis | Cell-level | Breast | 3 | 3 | - | ST Prediction |
| 2023-Dec | CRC-ST [28] | Molecular cartography uncovers evolutionary and microenvironmental dynamics in sporadic colorectal tumors | Spot-level | Bowel | 31 | 41 | 117,216 | ST Prediction, Clustering |
| 2024-Apr | SpatialTME [84] | The Web-Based Portal SpatialTME Integrates Histological Images with Single-Cell and Spatial Transcriptomics to Explore the Tumor Microenvironment | Spot-level | Multi-organ (12) | 26 cohort | 296 | - | Multi-tasks |
| 2024-Jun | HEST-1k [37] | HEST-1k: A Dataset for Spatial Transcriptomics and Histology Image Analysis | Multi-level | Multi-organ (26) | 153 cohort | 1,229 | 2.1M | Multi-tasks |
| 2024-Jun | STImage-1K4M [10] | STImage-1K4M: A histopathology image-gene expression dataset for spatial transcriptomics | Multi-level | Multi-tissue (50) | 121 cohort | 1,149 | 4.2M | Multi-tasks |
| 2025-Mar | Mouse-Brain-Xenium [60] | Optimizing Xenium In Situ data utility by quality assessment and best-practice analysis workflows | Cell-level | Brain | 1 | 4 | - | ST Prediction |
| 2025-May | OmiCLIP-ST [11] | A visual-omics foundation model to bridge histopathology with spatial transcriptomics | Multi-level | Multi-organ (32) | 113 cohort | - | 2.2M | Multi-tasks |
| 2025-Jun | CRC-HD [71] | High-definition spatial transcriptomic profiling of immune cell populations in colorectal cancer | Bin-level | Bowel | 5 | 5 | - | ST Prediction |

**TABLE III:**
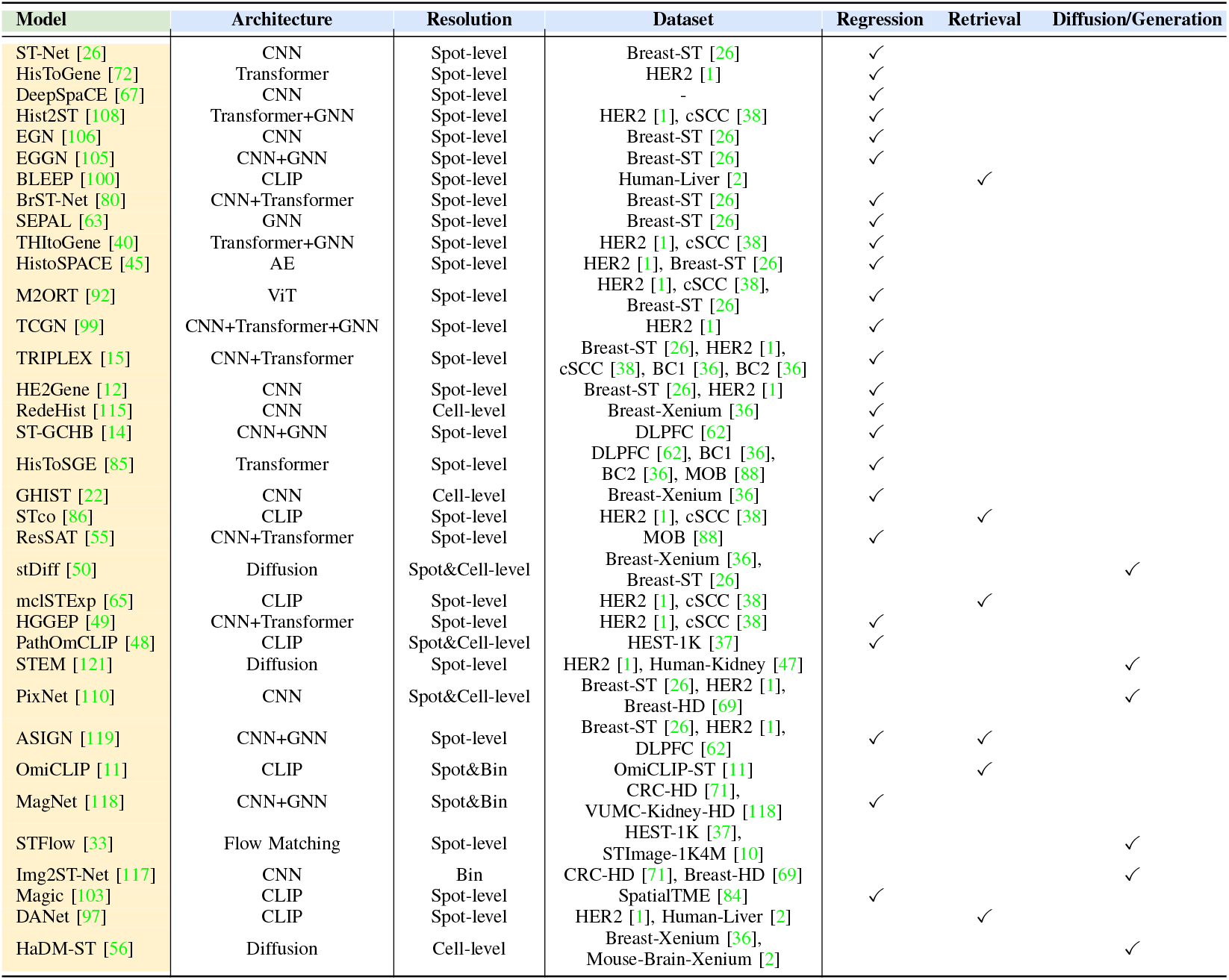
Comparison of collected methods for generative spatial omics task. The table summarizes the backbone architecture, output resolution, evaluated datasets, and the learning paradigm adopted for each model.

Early datasets were primarily generated at the spot level, where each capture area aggregates expression signals from multiple cells [1], [38], [47], [88]. Representative examples include the mouse olfactory bulb dataset [88], cSCC [38], and DLPFC [62]. These early resources typically contained only a limited number of samples, and thus mainly supported single-task analyses such as gene expression prediction. With the emergence of high-resolution platforms, ST research has gradually moved beyond coarse spot-level profiling toward finer bin-level and single-cell–level data. For example, Breast-HD [69] and CRC-HD [71] provide data at the bin level, while Breast-Xenium and Mouse-Brain-Xenium [60] achieve single-cell resolution, enabling more precise characterization of tissue microenvironments and cell–cell interactions.

Concurrently, the scale of ST datasets has expanded substantially. Early studies typically comprised only tens of thousands of spots with limited sample sizes. In contrast, modern resources now encompass millions of spots across over a hundred cohorts. Recent multi-level integrative datasets, such as HEST-1k [37], STImage-1K4M [10], and OmiCLIP-ST [11], consolidate multiple resolutions within a unified framework, further broadening both scale and coverage. For example, HEST-1k includes 153 cohorts with approximately 2.1 million spots, while STImage-1K4M spans 121 cohorts and about 4.2 million spots. Such large-scale expansion provides a stronger statistical foundation for robust analysis and establishes the conditions necessary for training large models.

Another notable trend is the expansion of organ coverage. Early datasets were often restricted to a single organ, such as the brain, breast, or skin, and were later extended to additional organs like the kidney and heart. More recently, large-scale resources such as SpatialTME [84] (covering 12 organs), HEST-1k (26 organs), and OmiCLIP-ST (32 organs) have achieved systematic multi-organ coverage. This shift enables studies to capture both cross-organ shared spatial patterns and organ-specific gene expression features.

Supported by these enriched datasets, the task paradigm has become increasingly diverse. What began as single-task efforts focused primarily on gene expression prediction has progressively expanded to include clustering, three-dimensional registration, and multi-task benchmarking. This evolution from single-to multi-task applications not only enhances the research value of the datasets but also drives methodological innovation.

## III. Vision-driven Tasks for Spatial Transcriptomics

Histopathological images are readily available at scale and capture rich morphological information that can be exploited for downstream ST analysis. The advancement of artificial intelligence and computer vision has provided powerful tools for medical image analysis [18], [44], [78], [120]. Recently, vision-driven ST models have emerged as a promising direction, offering a costeffective strategy to overcome experimental limitations and broaden data availability [26], [100], [119]. They have led a revolution in a wide range of downstream tasks in ST analysis [30], [67], [72], [106].

In this section, we introduce and review representative vision-driven tasks in spatial transcriptomics. We summarize how visual features extracted from histology images are leveraged to infer molecular landscapes, delineate spatial organization, and reveal functional heterogeneity within tissues.

### A. Generative Spatial Omics

Generative Spatial Omics establishes the foundation of the entire field. It leverages morphology-molecular correlations to directly infer gene expression profiles from histology images, thus offering a lightweight and cost-efficient strategy to overcome data broadening. Early models such as ST-Net [26] and HisToGene [72] were primarily designed with transfer learning strategies to improve gene expression prediction. Contrastive learning and multi-modelity learning then provided a new bridge to align vision-omics representation [65], [100]. With the emergence of larger datasets and more powerful models, recent approaches have further reinforced the central role of translation. For example, OmiCLIP [11] employs large-scale contrastive learning and multimodal pretraining to extend gene expression translation into a more universal and robust framework. By enabling costeffective reconstruction of molecular landscapes from tissue slides, translation provides the raw material for subsequent analyses and serves as a critical bridge between imaging and omics. A detailed illustration of learning paradigms and representative generative spatial omics frameworks will be presented in Section IV.

### B. Vision Feature Encoding and Clustering

Vision feature encoding serves as a fundamental step in vision-driven spatial transcriptomics, enabling models to transform histological textures and tissue architectures into informative embeddings. Clustering analysis then emerges as a key downstream task for revealing the spatial organization of cellular and molecular heterogeneity. The integration of image-derived embeddings allows for more accurate spatial alignment between histological morphology and molecular expression, thereby uncovering critical biological phenomena. Representative frameworks such as SpaGCN [30], SEPAL [63], and STAIG [104] incorporate visual features and graphbased modeling to enhance the characterization of local tissue structures and achieve precise identification of spatial domains. These methods collectively underscore the pivotal role of visual feature encoding in bridging histological organization with molecular expression patterns, establishing clustering as a cornerstone for spatial pattern discovery and tissue microarchitecture analysis [112], [114], [115].

### C. Super Resolution

Early ST platforms were constrained by low spot resolution, leaving substantial gaps between spots and limiting the availability of fine-grained spatial information. This limitation made it difficult to resolve spatial heterogeneity at the cellular or even subcellular level. To overcome this bottleneck, researchers have developed super-resolution prediction methods, which computationally refine spatial maps to higher granularity. By leveraging structural cues from histological images, models such as HiST2ST [108] and DeepSpaCE [67] can infer gene expression at sub-spot or even single-cell resolution, thereby markedly improving the accuracy and continuity of predicted molecular landscapes and enabling spatial analyses at cellular and subcellular scales.

At the same time, benchmarking efforts are increasingly shifting toward these new high-resolution settings. Methods such as MagNet [118] and PixNet [110] extend training and evaluation protocols to accommodate highresolution spatial transcriptomics data, further pushing the boundaries of model generalization and robustness.

### D. 3D Computer Vision Modeling

With the growing need to capture the full structural complexity of biological tissues [101], recent advances have introduced 3D modeling and reconstruction frameworks that integrate serial histological sections into volumetric molecular atlases. These models extend beyond the limitations of 2D analysis by explicitly modeling inter-slice spatial continuity and morphological consistency. Frameworks such as HisToSPACE [45] and STaCker [46] exemplify this paradigm. By aligning image and expression features across adjacent slices, they enable the reconstruction of organ-level spatial context and continuous molecular landscapes.

For other aspects, ASIGN [119] further leverages 3D spatial relationships and introduces a new learning paradigm that transitions from conventional 2D WSI-ST prediction to partially observed 3D volumetric WSI-ST imputation. Such 3D alignment and modeling substantially broadens the analytical scope of spatial transcriptomics. 3D computer vision modeling is emerging as a critical frontier in vision-driven spatial omics, bridging micro-scale morphology with macro-scale biological context.

## IV. Learning Manners for Generative Spatial Omics

Generative spatial omics is one of the most common and widely studied tasks in vision-driven models. Existing approaches can be broadly categorized into three types: (1) regression-based prediction methods, which directly fit the mapping between image features and gene expression levels through end-to-end supervised learning; (2) retrieval-based contrastive learning methods, which align image and expression spaces and perform prediction by retrieving similar expression patterns from large-scale databases; and (3) diffusion-based generative frameworks, which employ generative modeling to reconstruct high-dimensional expression distributions and enable richer molecular-level expression generation during inference.

Figure 3 summarizes the typical workflows of these three approaches in both training and inference. As shown, each strategy differs in its modeling perspective, training objective, and prediction output. In this section, we provide a systematic overview of these mainstream strategies and further discuss existing methods that extend or build upon them.

**Fig. 3:**
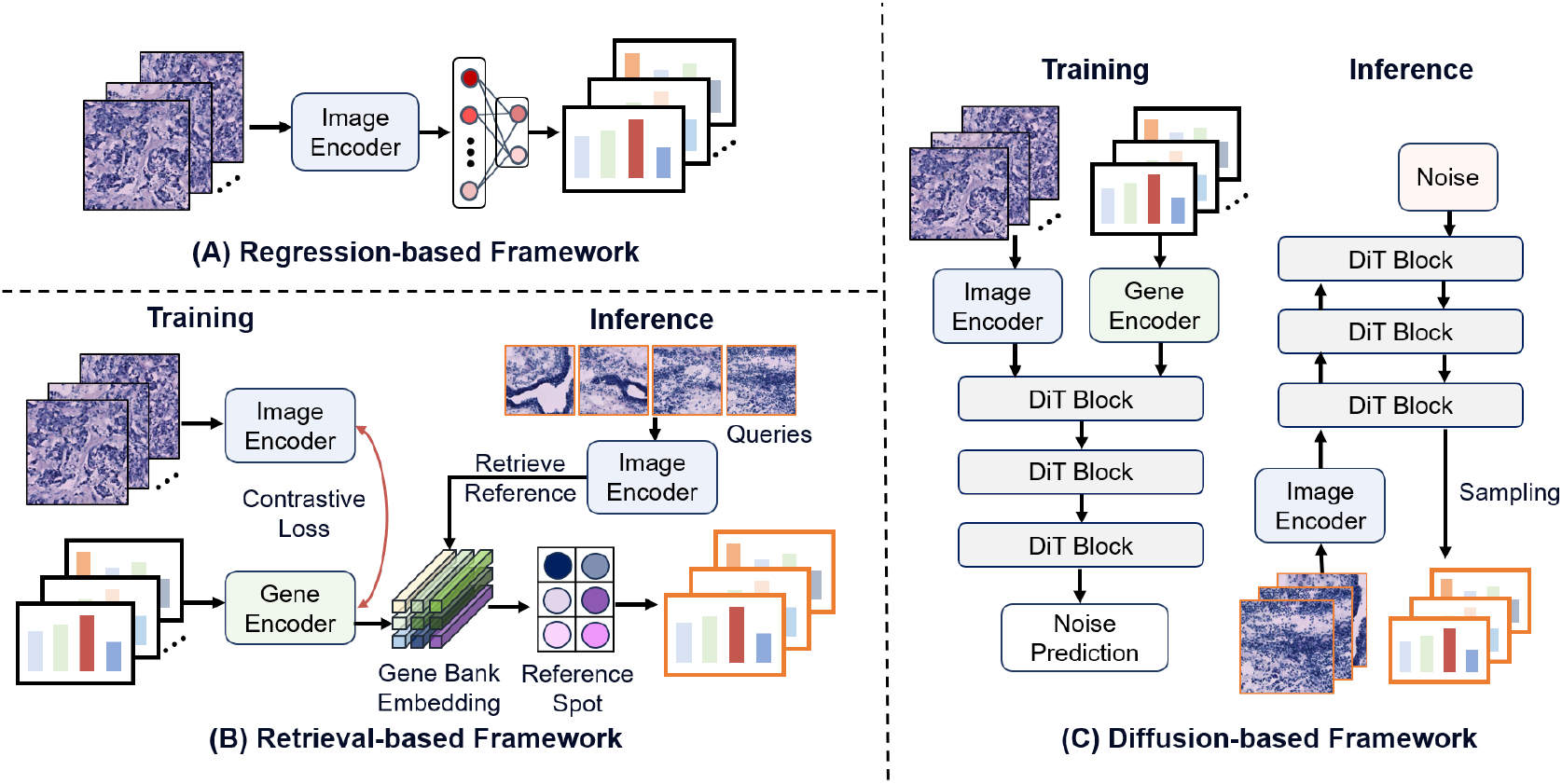
Learning paradigms for generative spatial omics Task. Current learning paradigms mainly include three forms: (A) Regression-based prediction which directly maps histology tiles to gene expression profiles; (B) Retrieval-based searching that aligns image–gene representations and retrieves nearest reference spots to form predictions; (C) Diffusion-based generation which performs conditional noise prediction with a DiT and, at inference, denoises from noise to generate expression maps.

### A. Regression-based Modeling

Regression-based approaches are among the earliest and most widely adopted strategies for Image-to-ST translation. They formulate gene expression prediction as a multi-dimensional regression task, where features extracted from histology images are mapped into the gene expression space through regression layers. An early representative work, ST-Net [26], first introduced this idea. Leveraging transfer learning, ST-Net employs a pre-trained CNN backbone [32] to extract image features, which are then passed through linear layers to project them into the dimensionality of gene expression. Formally, given a histology image *I* with feature representation *f* (*I*), the predicted gene expression vector *ŷ* can be expressed as:

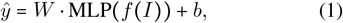

where *W* and *b* denote the weights and bias of the regression layer, respectively.

The regression-based training method offers a simple and efficient solution under end-to-end supervision. Nonetheless, its formulation as independent spot-wise regressions restricts the modeling of spatial dependencies, while its predictive performance is bottlenecked on the quality of extracted features. To address these limitations, subsequent studies have proposed a variety of strategies to enhance spatial association modeling [39], [72], [108], [119] and semantic feature representation [99], [105], [106].

HisToGene [72] and His2ST [108] first utilize spatial coordinates to construct connections between spots, thereby refining the spatial representation of ST data through contextual information from neighboring regions. EGN [106] and EGGN [105] then adopt image similarity as a metric for constructing connections, bringing spots with similar visual tissue characteristics closer in the feature space to better guide expression prediction. ASIGN [119] further extends modeling into 3D space by combining spatial relationships and feature similarity, extrapolating known 2D label information across tissue layers to achieve more accurate 3D ST data prediction. In addition, TCGN [99] incorporates diverse backbone architectures in the feature extraction stage, combining the strengths of CNNs, GNNs, and Transformers to comprehensively capture local patterns, spatial relationships, and long-range dependencies.

### B. Retrieval-based Alignment

Current success of Contrastive Language–Image Pretraining (CLIP) in vision–language modeling [79] has opened new possibilities for joint image–ST representation learning. ST is a biological language, where gene expression profiles serve as semantic tokens describing the molecular state of tissue [95]. This perspective motivates the adaptation of contrastive learning frameworks to align histology images with transcriptomic signals, thereby enabling more effective cross-modal representation and downstream prediction [11], [65], [100], [103]. BLEEP [100] introduced the first multi-modal framework, where two encoders are used to map images and ST gene expression vectors into a shared latent feature space. The core idea is to maximize the similarity between an image and its paired gene expression vector while minimizing the similarity with mismatched pairs. During the inference stage, gene expression prediction is achieved by retrieving the most similar known spots.

Formally, for the *i*-th spot consisting of an image embeddings are obtained through an image encoder *f* (·) *I*_*i*_ and its corresponding gene expression vector *x*_*i*_, the and a gene expression encoder *g*(·):

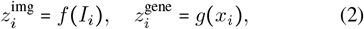

The contrastive learning objective is based on a symmetric InfoNCE loss, which jointly optimizes image-to-gene and gene-to-image alignment:

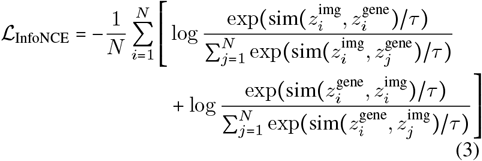

where sim, denotes cosine similarity and *τ* is a temperature parameter.

During inference, BLEEP predicts gene expression by comparing the embedding of a query image with all gene expression embeddings in a reference database. Given a query image *I*_*q*_ with embedding 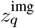, the similarity scores with all database embeddings 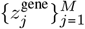 are com-puted as:

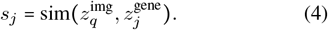

The top-*K* most relevant gene expression vectors 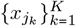 are then retrieved, and the final prediction is obtained by a weighted aggregation:

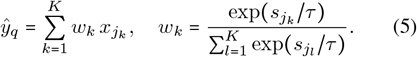

Building on this framework, mclSTExp [65] extends BLEEP by incorporating spatial coordinates to enable joint spatial modeling, thereby better capturing the spatial dependencies between histology images and transcriptomic signals.

Meanwhile, with the growing scale of ST data, crossmodal alignment tasks have expanded from the original image–gene expression setting to richer modalities such as image–ST gene sentence, where sets of genes or semantically enriched gene descriptions serve as alignment targets. Such extensions not only enhance the semantic expressiveness of the representation space, but also provide new opportunities for the development of general vision–omics foundation models [11].

### C. Generation-based Paradigm

The generation-based paradigm attempts to directly model the joint distribution between histology images and gene expression. In this way, ST data can be obtained through a generative mechanism during inference [33], [45], [117], [121]. In recent years, the emergence of diffusion models [29] has provided new opportunities for this task.

STEM [121] first incorporates histology images as conditional variables, using DiT [73] as the backbone to reconstruct gene expression matrices in a generative manner. By modeling the probabilistic generation process, STEM captures complex distributions and provides more stable and diverse outputs than conventional regression-based approaches. Formally, let a gene expression vector be *x*_0_ ∈ ℝ^*G*^, where *G* denotes the number of genes. The forward diffusion process is defined as:

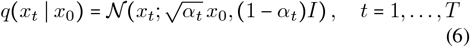

where *x*_*t*_ denotes the noisy sample at step *t*, and 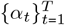 is the noise schedule.

In the reverse generation process, the model learns the conditional distribution:

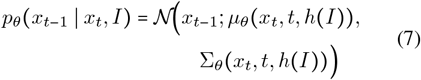

where *I* denotes the histology image and *h I* is the conditional embedding extracted by an image encoder (e.g., a DiT backbone). Training is typically performed by minimizing the noise prediction objective, equivalent to a variational lower bound:

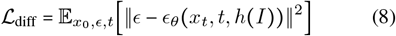

where *ϵ* ∼ 𝒩 ( 0, *I*) denotes Gaussian noise, and *ϵ*_*θ*_ is the noise predicted by the model.

During inference, given a histology image *I* as a condition, the diffusion model starts from Gaussian noise *x*_*T*_ ∼ 𝒩 ( 0, *I*) and progressively applies the conditional denoising process to generate the final gene expression prediction 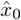.

More recently, STFlow [33] introduces Flow Matching [54], a novel generative modeling framework that directly learns a deterministic, time-dependent vector field *v*_*θ*_(*x, t*). STFlow uses histology images as conditional inputs, where the conditional embedding *h*(*I*) guides the deterministic transport from a Gaussian prior to the gene expression distribution:

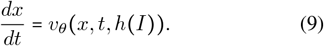

Thus, flow Matching reduces the number of sampling steps and improves both the stability and diversity of the generated results.

Overall, the existing regression-based, retrieval-based, and generative methods provide diverse strategies and perspectives for image-to-ST translation. It is important to note, however, that generative spatial omics is not confined to these paradigms. With the continuous accumulation of large-scale ST data and the rapid development of multimodal learning techniques, new methods and frameworks are expected to emerge, further expanding the boundaries and potential of this task.

## V. Empirical Implementation and Evaluation Metrics

Current vision-driven models lack a unified training pipeline or standardized preprocessing framework. Variations in data preprocessing, gene selection, and training and validation schemes lead to diverse experimental implementations. In this section, we review existing models from these three aspects and summarize the commonly used evaluation metrics. Table IV presents the implementation strategies of existing methods and their performance on specific datasets.

**TABLE IV:**
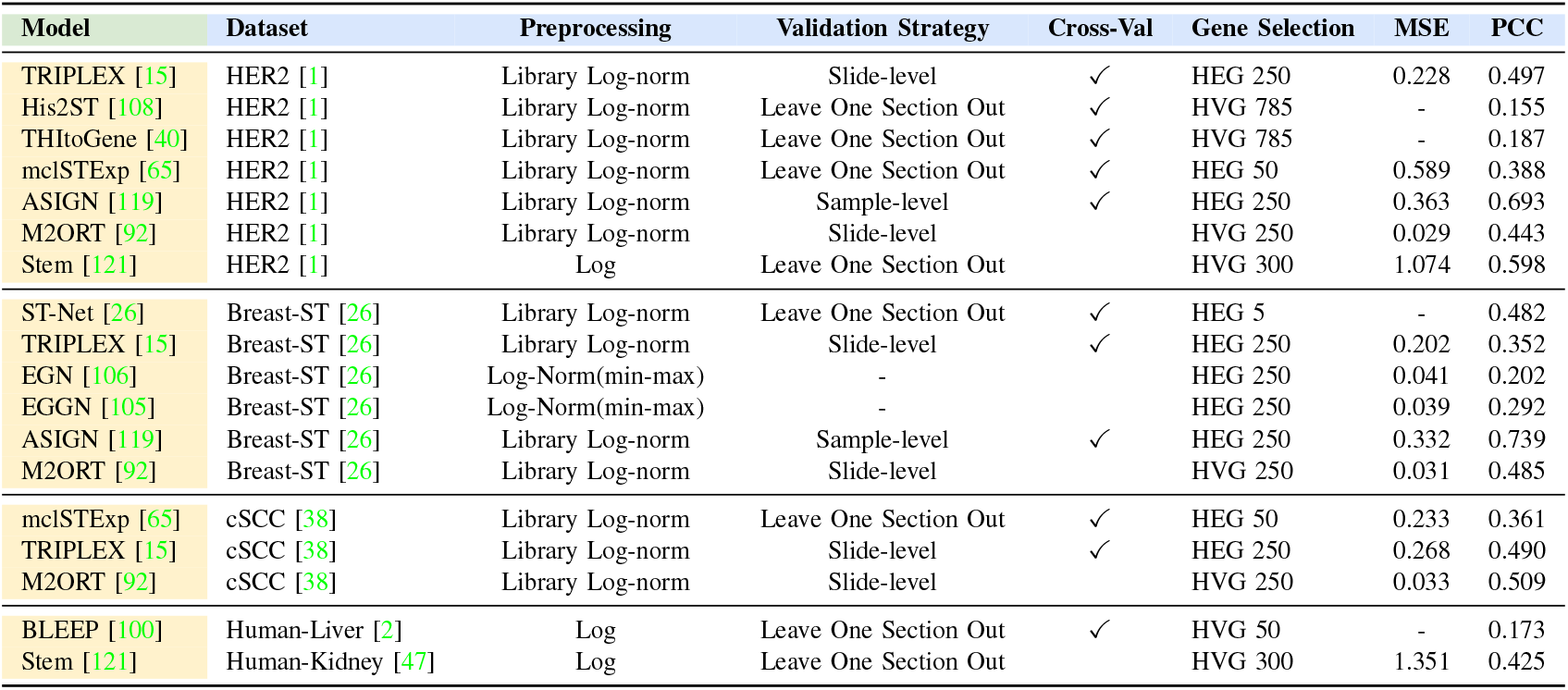
Comparison of vision-to-ST models across datasets, preprocessing pipelines, validation strategies, and performance. All metrics are taken directly from the original papers.

### A. Implementation for Generative Spatial Omics

#### 1) Gene Selection

Gene selection is the starting point of experimental design and a critical factor influencing both model performance and generalization. Due to the inherent sparsity of gene expression in ST data, early studies [26], [106], [119] commonly adopted the strategy of selecting the highest expression genes (HEGs) as prediction targets [26]. However, relying solely on highly expressed genes may overlook important spatial information. To better capture the correspondence between local tissue structures and molecular states, many methods [100], [121] have therefore proposed using highly variable genes (HVGs) as training targets, thereby enhancing the model’s sensitivity to spatial heterogeneity. In addition, there are substantial differences in the scale of gene selection across methods. Some studies [65], [100] focus on a relatively small set of key genes (e.g., 50 targets), emphasizing specific molecular pathways or downstream tasks, whereas others [39], [108] extend the selection to encompass hundreds of genes from the entire dataset (up to 785 genes), aiming to cover the transcriptomic space as comprehensively as possible. The diversity of gene selection strategies reflects different trade-offs between biological interpretability and computational complexity, and these choices directly shape the training and validation schemes.

#### 2) Preprocessing

Existing gene expression preprocessing methods can generally be divided into two categories. The first applies a direct logarithmic transformation to the raw counts in order to mitigate the long-tail distribution effect commonly observed in ST data [100], [121], thereby smoothing the numerical differences between lowly and highly expressed genes. The second approach normalizes the raw counts first and then log-transforms the normalized values [15], [26], [92], [105], [119].

In practice, the second approach has gradually become the mainstream choice. This method not only reduces the influence of extreme values but also effectively alleviates the bias introduced by differences in sequencing depth. It is typically implemented as a combination of Library size normalization and log transformation, also referred to as Library Log-norm. Specifically, let *x*_*i,j*_ denote the raw expression of gene *j* in spot *i*. The Library size normalization is performed as:

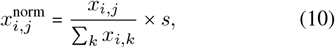

where ∑_*k*_ *x*_*i,k*_ represents the total expression (library size) of spot *i*, and *s* is a scaling factor (commonly 10^4^ or 10^6^). Subsequently, a log transformation is applied:

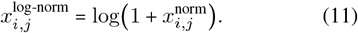

This strategy ensures the comparability of expression values across different spots, compresses the dynamic range of the distribution, and provides more stable and robust input features for subsequent model training.

In addition to conventional numerical preprocessing, some methods [17], [91] further discretize the raw counts into bins to mitigate differences in gene expression across cohorts. This binning strategy reduces data sparsity and noise, while also providing a standardized input representation for training large-scale foundation models.

#### 3) Validation Strategy

Different methods show substantial variation in the split of ST data. Early studies [15], [26] commonly adopted a slide-level split for cross-validation, where different sections from the same sample were treated as independent training and validation units. Building on this, another widely used approach [39], [92], [100] is Leave-One-Section-Out, where one section is held out for validation while the others are used for training.

However, ST data are often derived from multiple layers or adjacent regions of the same sample, and these sections usually share strong similarities in both tissue morphology and molecular distribution, thereby inflating the reported model performance. To address this issue, recent studies have proposed more stringent splitting strategies. For instance, some methods [119] adopt a sample-level split, ensuring that all sections from the same sample are confined to either the training set or the validation set. Others [15] further employ external validation, where models are evaluated on entirely independent datasets, providing a more objective measure of generalization ability.

Notably, recent studies [90] have increasingly recognized the disparities in the experimental implementation of existing methods and have begun to conduct systematic benchmarking under more unified and fair experimental conditions. Such efforts enable a more objective evaluation of model performance within comparable settings and drive the field toward a more transparent and reproducible research paradigm.

### B. Evaluation Metrics

To systematically evaluate the performance of Image-to-ST models, we summarize the commonly used metrics in existing studies in Table V. According to their design purposes and application scenarios, these metrics can be broadly categorized into error-based, correlation-based, variation-based, and clustering-based.

**TABLE V:**
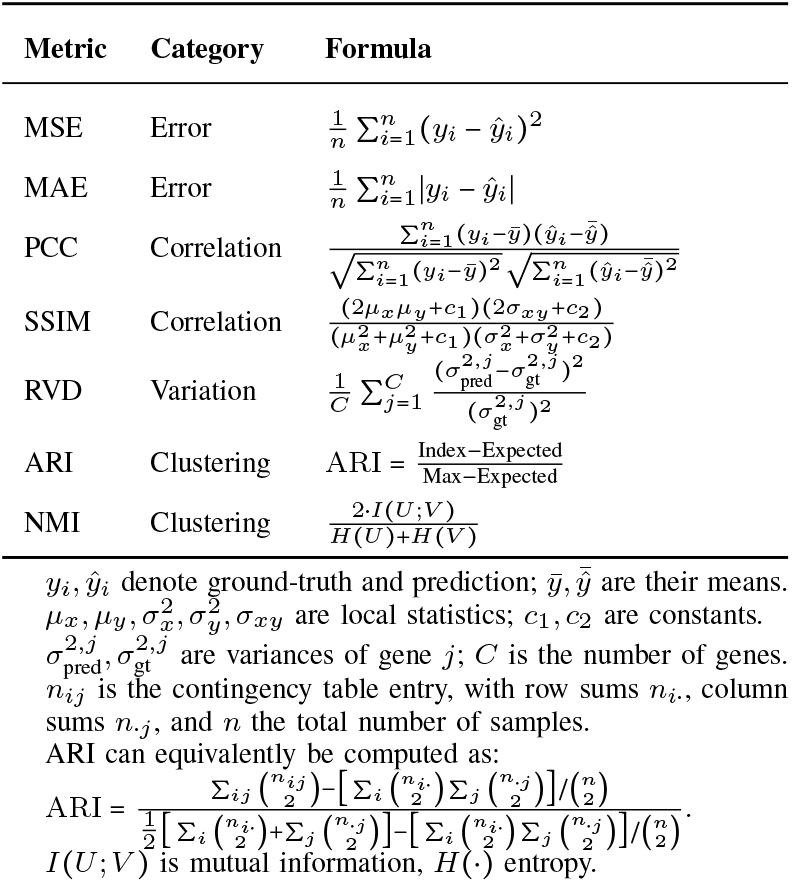
Common evaluation metrics used in vision-to-ST tasks. The metrics are grouped by category with their formulas.

#### 1) Error-based Metrics

Mean Squared Error (MSE) and Mean Absolute Error (MAE) are the most widely used metrics in ST prediction [15], [26], [119], mainly for regression tasks that aim to directly predict gene expression levels. MSE is more sensitive to large deviations, highlighting the impact of extreme errors, while MAE is more robust to outliers and provides a balanced estimate of average error. They quantify the numerical discrepancy between predictions and ground truth, serving as core measures of predictive accuracy.

#### 2) Correlation-based Metrics

The Pearson Correlation Coefficient (PCC) measures the linear correlation between predicted and ground-truth expression profiles and is a key indicator of whether models can capture spatial trends of gene expression [26], [39], [99]. However, in high-resolution and sparse data, PCC can be strongly affected by excessive zero values, reducing its reliability.

In contrast, the Structural Similarity Index (SSIM), originally designed for image quality assessment, has been adopted in ST tasks to evaluate whether predicted spatial maps preserve structural patterns consistent with tissue morphology [49], [102], [117]. By jointly considering local means, variances, and covariance, SSIM provides a more robust measure of spatial similarity, particularly under sparse data conditions.

#### 3) Variation-based Metrics

Relative Variation Distance (RVD) measures the discrepancy between predicted and ground-truth variance structures at the gene level [121]. Prior studies have noted that models may achieve high PCC scores even when merely outputting mean predictions with little variation, thereby masking true spatial differences [100], [121]. Therefore, RVD is proposed to tell whether a model can faithfully capture the biological heterogeneity inherent in spatial transcriptomics data. Thus, RVD serves as a complementary metric to existing evaluation systems, helping to identify models that perform well on correlation metrics but fail to capture heterogeneity.

#### 4) Clustering-based Metrics

Adjusted Rand Index (ARI) and Normalized Mutual Information (NMI) are commonly used clustering evaluation metrics, primarily applied to downstream tasks such as spatial domain identification [9], [30], [74]. ARI measures the consistency between predicted and ground-truth cluster assignments while correcting for random agreement, whereas NMI evaluates the amount of shared information between predicted and true partitions from an information-theoretic perspective. Thus, these metrics reflect the model’s ability to capture cellular heterogeneity and cell niches..

## VI. Discussion

While substantial progress has been achieved in data resources and modeling paradigms, several key challenges remain, including data heterogeneity, limited reproducibility, and the lack of biologically grounded evaluation standards. In this section, we discuss these pressing issues and further outline potential learning strategies and translational opportunities, providing a roadmap for the next phase of vision–omics integration.

### A. Challenges and Future Directions

#### 1) Data Integration and Processing

A primary challenge for vision-driven ST models lies at the data level. Training high-quality deep learning models depends on abundant, high-quality ST data, yet such data remain scarce in practice. Although spatial transcriptomics platforms have advanced rapidly and increased data throughput, sample sizes across diverse settings are still insufficient to support large-scale training and robust generalization. In addition, substantial heterogeneity exists across sequencing platforms in capture efficiency and sequencing depth [60]. These differences manifest in data formats and signal strength, greatly complicating cross-platform comparison and model transfer [24].

Compounding matters, histopathology images and transcriptomic signals cannot always be perfectly aligned in space. Such spatial misalignment may arise from section deformation or tissue cutting offsets [25], [58]. Systematic biases, including batch effects, variations in experimental processing, and differences in sample preparation, can further obscure biological signals or introduce spurious correlations. In many cases, conventional library-size normalization can even impair the delineation of spatial domains [6].

Substantive progress will require breakthroughs in data integration and standardization. On one hand, the community should accelerate the construction of larger, more balanced resources [10], [37] to mitigate distributional bias in training data. On the other hand, establishing unified preprocessing and analysis pipelines [31], [64], including cross-platform standardization and batcheffect correction, is essential to ensure comparability and consistency across sources. In parallel, designing robust models that maintain stable performance under heterogeneous inputs will be key to improving the practicality and deployability of vision-driven ST models.

#### 2) Toward Fair and Reproducible Validation

Existing studies lack unified and fair training–validation protocols. First, data partitioning is inconsistent. Many works adopt leave-one-section-out [39], [100] or slide-level random splits [15], [92], which often place samples from the same patient/tissue, which shares highly correlated spatial and morphological characteristics in the training and test sets, causing information leakage and systematically inflating performance. In contrast, splits at the patient or center level better reflect clinically meaningful external generalization [119].

Second, most methods report results on a single dataset only, with limited cross-dataset and crossplatform evaluation [15]. As a result, model robustness to distribution shift, batch effects, and experimental variation remains unclear. Finally, discrepancies in test-gene sets and preprocessing pipelines are another major source of bias. Differences in gene selection, normalization, and quality control can substantially alter reported metrics and thus distort horizontal comparisons.

Recent efforts have begun to establish more equitable and reproducible evaluation frameworks to systematically assess method performance [90]. On one hand, the community should build cross-cohort benchmarks [10], [37] and standardize partition schemes, gene selection, and preprocessing to avoid performance artifacts stemming from split and pipeline choices. On the other hand, open-sourcing code and weights, locking dependencies and random seeds, and providing one-click evaluation entries will improve reproducibility and transparency. Through such normalization and openness, the field of vision models can move beyond “works on this dataset” toward cross-dataset, cross-platform, and deployment-ready standards.

#### 3) Exploration of Evaluation Metrics

The limitations of current evaluation protocols constitute a major bottleneck. Most studies still rely on numerical error metrics (e.g., MSE, MAE) to assess prediction accuracy. However, these primarily reflect curve-fitting capability and are insufficient to determine whether models truly capture biologically meaningful spatial patterns. Meanwhile, traditional correlation measures (e.g., PCC) can break down on highly sparse datasets [117] or under “mean-like” predictions [121], yielding spuriously high or low correlations that mislead model assessment.

Looking ahead, evaluation can advance along two axes: multidimensionality and standardization. Beyond retaining regression-style error metrics, we should incorporate spatial and biological criteria, such as markergene recovery, to directly reflect biological interpretability and spatial fidelity. In parallel, we advocate developing mechanism-oriented metrics, including pathway-level recovery and the detectability of disease-relevant markers. Such metrics help shift the focus from mere “numerical fit” to “mechanistic concordance,” enabling a more comprehensive appraisal of a model’s true value for biological interpretation and clinical application.

#### 4) Applications and Potential Learning Strategies

Despite the broadening of task scope, existing vision-driven ST studies still focus on foundational objectives such as gene expression prediction and spatial clustering, with limited progress toward clinical translation. Bridging basic research and clinical practice, therefore, remains a pressing challenge [35], [87]. At the same time, current models lack a unified framework for multimodal integration, making it difficult to jointly incorporate histopathology, transcriptomics, proteomics, and other omics to form a more comprehensive and interpretable tissue representation [11].

Looking ahead, progress can proceed along two main fronts. First, develop cross-modality alignment and fusion strategies, such as joint embedding spaces and generative modeling, to move toward a unified vision–omics representation. Second, adopt efficient training paradigms to alleviate the computational burden, including self-supervised learning, cross-modal pretraining, and parameter-efficient tuning. Coupled with sustained collaboration between clinicians and domain experts, these advances will improve scalability and adaptability, expediting deployment in real-world applications such as precision medicine and early disease screening.

### B. Limitations

This review concentrates on vision-driven models for spatial transcriptomics analysis published between 2020 and September 2025. Although we aimed to cover the most representative advances, the selection is inevitably constrained by publication timing and the availability of public evaluations. As a result, some very recent contributions, particularly those appearing near the end of this window or lacking thorough benchmarking, may not be included. Readers are therefore encouraged to consult the latest releases and datasets beyond this survey to complement our coverage.

It is also worth noting that both computer vision and spatial transcriptomics are evolving at high speed, with new techniques and modeling paradigms continually reshaping the landscape. Maintaining an accurate and complete picture will require ongoing literature tracking and periodic updates. Accordingly, this article should be viewed as a time-bounded overview rather than a definitive account. To keep pace with emerging work and trends, subsequent studies and new evidence should be monitored and incorporated as they appear.

## VII. Conclusion

In this comprehensive survey, we have systematically reviewed representative advances in vision-driven models for ST analysis, categorizing existing approaches from multiple perspectives, including model architectures, learning paradigms, downstream tasks, and data resources. vision-driven ST models are moving beyond the limitations of regression-based paradigms toward multi-model and generative methods. Their applications have also expanded from gene expression prediction and clustering to more complex endeavors like 3D reconstruction. This shift reflects both methodological diversification and the vast potential of these approaches in practical applications. Nevertheless, opportunities come with challenges. Current research remains constrained by the scarcity and heterogeneity of high-quality datasets, the absence of standardized and fair evaluation protocols, and limited cross-platform generalization. These issues underscore that vision-driven ST is still in a phase of rapid evolution. Looking ahead, advancing this field will require more efficient model architectures, deeper integration of multi-modal information, and the construction of larger, high-quality benchmark datasets. In conclusion, the evolution of vision-driven ST models represents a new frontier in the integration of bioinformatics and medical imaging. Beyond addressing the practical limitations of experimental technologies, these models open new pathways for understanding complex tissue organization and disease mechanisms, positioning themselves as a vital bridge between basic research and clinical application.

## Acknowledgements

This research was supported by NIH R01DK135597 (Huo), DoD HT9425-23-1-0003 (HCY), and KPMP Glue Grant. This work was also supported by Vanderbilt Seed Success Grant, Vanderbilt Discovery Grant, and VISE Seed Grant. This project was supported by The Leona M. and Harry B. Helmsley Charitable Trust grant G-1903-03793 and G-2103-05128. This research was also supported by NIH grants R01EB033385, R01DK132338, REB017230, R01MH125931, and NSF 2040462. We extend gratitude to NVIDIA for their support by means of the NVIDIA hardware grant. This work was also supported by NSF NAIRR Pilot Award NAIRR240055.

